# Reconstituted high-density lipoproteins rescue diabetes-impaired endothelial cell metabolic reprograming and angiogenic responses to hypoxia

**DOI:** 10.1101/2024.08.16.608361

**Authors:** Khalia R. Primer, Joanne T.M. Tan, Lauren Sandeman, Victoria A. Nankivell, Liam G. Stretton, Emma L. Solly, Peter J. Psaltis, Christina A. Bursill

## Abstract

**Objective:** Impaired angiogenic responses to ischemia underlie diabetic vascular complications. Reconstituted high-density lipoproteins (rHDL) have proangiogenic effects in diabetes. The pyruvate dehydrogenase kinase 4 (PDK4)/pyruvate dehydrogenase complex (PDC) axis is an oxygen-conserving mechanism that preserves EC functions in hypoxia. We aimed to determine the role of the PDK4/PDC axis in angiogenesis, the effect of diabetes on its regulation in response to ischemia, and in the proangiogenic properties of rHDL.

**Approach and Results:** In a murine wound healing model, PDK4 and pPDC were elevated early (24h) post induction of wound ischemia in non-diabetic wounds, which did not occur in diabetic mice. Topical rHDL rescued this impairment, enhancing PDK4 (68%, P<0.05) and pPDC (165%, P<0.01) in diabetic wounds. In parallel, wound neovascularization (62%, P<0.05) and closure (154%, P<0.0001) were increased in diabetic rHDL-treated wounds. In vitro, PDK4 and pPDC levels were increased in ECs exposed to hypoxia (65%, 70% respectively, P<0.05). High glucose did not elicit a further step-wise induction in PDK4/pPDC, with aberrant increases in mitochondrial respiration (19%, P<0.05), coupled with impaired EC angiogenic functions. Importantly, rHDL increased PDK4 and pPDC two-fold, returning mitochondrial respiration and EC angiogenic functions to normal glucose levels. In vitro, PDK4 siRNA knockdown attenuated the proangiogenic effects of rHDL. In vivo PDK4 inhibition ameliorated topical rHDL-mediated increases in wound angiogenesis and closure. Using chromatin immunoprecipitation, rHDL increased forkhead box O1 (FOXO1) binding to the PDK4 promoter and suppressed FOXO1 phosphorylation, presenting FOXO1 as a mechanism for the induction of PDK4 by rHDL.

**Conclusion:** The PDK4/PDC axis response to ischemia is impaired in diabetes and important for the proangiogenic effects of rHDL.

## INTRODUCTION

Diabetes mellitus (DM) is an increasing global epidemic and poses a significant health and economic burden.^1,2^ The vascular complications of DM have a major impact on a patient’s quality of life and are strongly associated with impaired angiogenic responses to ischemia. Patients with DM exhibit reduced angiogenesis in response to the tissue ischemia caused by athero-occlusions in femoral and coronary arteries.^3^ Cutaneous wound healing is also impaired in DM as angiogenic responses to wound ischemia are essential for healing.^4^ Patients with DM therefore have a higher risk of lower-limb amputation than their non-diabetic counterparts.^2,4,5^

Angiogenesis is the process by which ECs respond to tissue ischemia by forming new blood vessels from pre-existing ones. Cellular metabolic adaptation to hypoxia is a key aspect of this response,^6^ wherein ECs must reduce their oxygen consumption to conserve energy and avoid oxidative stress.^7,8^ To achieve this, ECs reprogram their metabolism to increase anaerobic glycolysis^9^ and reduce mitochondrial respiration.^10^ An inability to undergo this switch in hypoxia may underpin inadequate neovascularization in DM.

A mechanism which regulates this switch is the mitochondrial pyruvate dehydrogenase kinase 4 (PDK4)/pyruvate dehydrogenase complex (PDC) axis.^11^ In response to hypoxia, PDK4 phosphorylates the E1α subunit of the PDC to inactivate it.^12,13^ The elevation of PDK4 in hypoxia therefore decreases the amount of glucose-derived acetyl CoA available to fuel the tricarboxylic acid (TCA) cycle, increasing metabolic efficiency in hypoxia and preserving cell survival, which is essential for EC participation in angiogenesis.^6^ The role of the PDK4/PDC axis in EC angiogenesis and the effect of diabetes is relatively unexplored.

Reconstituted high-density lipoproteins (rHDL) have been shown to rescue high glucose-impaired angiogenesis *in vitro* and *in vivo* in diabetic models.^14^ However, its ability to correct EC metabolic responses to hypoxia in diabetes is unknown. Accordingly, the objectives of this study were to investigate the importance of the PDK4/PDC axis in diabetes-impaired angiogenesis *in vivo* in a murine model of diabetic wound healing and in ECs *in vitro*, and determine the ability of rHDL to correct this pathway.

## MATERIALS AND METHODS

### Isolation of apoA-I

Isolated human plasma HDLs were isolated from pooled samples of 5 normal donors, for which the HDL fraction (1.063–1.21g/mL) was separated by ultracentrifugation and then dialyzed against phosphate-buffered saline (PBS). ApoA-I was isolated from lipid free HDL by ultracentrifugation and anion exchange chromatography on a 2.6x24.0cm Q Sepharose Fast Flow column (Amersham Pharmacia Biotech) attached to an AKTA Fast Performance Liquid Chromatography (FPLC) system (Amersham Pharmacia Biotech) as previously described.^15^ Delipidated HDL was dissolved in 20mM Tris, 6M urea and filtered before being loaded onto the column. ApoA-I was separated and eluted from the column using an increasing gradient of 20 mM Tris, 0.2M NaCl, 6 M Urea (pH 8.5). Protein elution was monitored at A280 nm and the fractions corresponding to the single peak for apoA-I were pooled. ApoA-I was extensively dialyzed against 20 mM ammonium bicarbonate, lyophilized and stored at −20°C.

### Preparation of apoA-I, phospholipid vesicles, discoidal rHDL, glycated rHDL and oxidized rHDL

Lipid-free apolipoprotein A-I (apoA-I) was reconstituted in buffer (100mM Tris, 3M Guanidine HCl, pH 8.2), and dialyzed against Tris-buffered saline (TBS). 1-palmitoyl-2-linoleoyl-phosphatidylcholine (PLPC) vesicles were prepared by sonication (3 cycles of 5min each on ice) in PBS and cholate and dialyzed against PBS. Discoidal reconstituted HDL (rHDL) was prepared by complexing apoA-I with PLPC at an initial PLPC/apoA-I molar ratio of 100:1 (final PLPC/apoA-I molar ratio is approximately 80:1). rHDL was dialyzed against TBS and PBS. Glycated rHDL (Glyc-rHDL) was prepared by incubating discoidal rHDL with 3.0mmol/L methylglyoxal for 24h at 37°C under 5% (v/v) CO_2_.^16^ Unreacted methylglyoxal was removed by dialysis against PBS. Oxidized rHDL (Ox-rHDL) was prepared by incubating discoidal rHDL (final apoA-I concentration 1mg/mL) with an equal volume of 50mmol/L PBS (pH 7.4) containing 0.2mmol/L hydrogen peroxide, 0.2 mmol/L diethylene triamine pentaacetic acid, 0.4 mmol/L L-tyrosine, and 20 nmol/L myeloperoxidase at 37°C for 24h under 5% (v/v) CO_2_, then dialyzed against PBS.^17^ rHDL, Glyc-rHDL and Ox-rHDL were filtered with a 0.22μm low protein-binding low-retention filter (Merck Millipore, Ltd.). Protein concentrations of apoA-I, rHDL, Glyc-rHDL and Ox-rHDL were determined using a Pierce Bicinchoninic Acid (BCA) Protein Assay Kit (Life Technologies) and phospholipid concentration was determined enzymatically (Wako). rHDL, Glyc-rHDL and Ox-rHDL were run on a homogeneous 20% SDS-polyacrylamide gel (Life Technologies) with Coomassie blue staining to show purity and size distributions (Supplementary Figure 1A). rHDL size distribution was also determined using dynamic light scattering (mean diameter: 10.69 nm; Supplementary Figure 1B). rHDL was stored under N_2_ gas and tubes sealed with paraffin for up to two weeks at 4°C.^14^

### Animal Studies

All procedures were conducted with ethical approval from the SAHMRI Animal Ethics Committee (#SAM301) and conformed to the Guide for the Care and Use of Laboratory Animals (NIH). In specific pathogen free housing in cages of four, male 6-week-old C57Bl/6J mice (Jackson Laboratory, USA) were rendered diabetic two weeks prior to surgery by a bolus intraperitoneal injection of streptozotocin (165µg/g). A parallel study was conducted in which female 6-week-old C57Bl/6J mice (Jackson Laboratory, USA) were rendered diabetic three weeks prior to surgery by a bolus intraperitoneal injection of streptozotocin (180µg/g). One-week post-injection, hyperglycaemia was confirmed using the Accu-CHEK Performa Blood Glucometer. A blood glucose level of 15.0mmol/L or above was considered diabetic.

### Murine Wound Healing Model

The wound healing model was conducted as previously described.^14,18^ Briefly, under inhalation isoflurane (3%) anaesthesia, two full thickness wounds were placed on the back flanks and secured with silicon splints (Sigma), attached with interrupted sutures. For each mouse, one wound received topical rHDL (50μg/wound/day) or PBS (vehicle control), applied daily. A dressing (Opsite^TM^) was applied, and digital images and wound area measurements were taken daily. Wound blood perfusion was determined using laser Doppler imaging. After completion of each time point, mice were humanely killed using an overdose of isoflurane anaesthesia (5%), followed by cardiac puncture, and the wound area was excised.

### Immunohistochemistry

The excised wound area was cut in half and placed in 30% sucrose for 24 hours, followed by 4% paraformaldehyde for 24 hours, then in 70% ethanol at 4°C. In a tissue processor, wound tissue was infused with paraffin using a sequence of graded ethanol concentrations, xylene and then paraffin in a 6-hour cycle. The wounds were then paraffin-embedded and 5μm sections were taken and mounted on SuperFrost+ slides (Thermo Scientific). Prior to staining, slides were deparaffinised using xylene, then rehydrated in decreasing ethanol concentrations.

For antigen retrieval, slides were boiled for 20min in sodium citrate buffer (10mM sodium citrate, 0.05% Tween 20, pH 6.0). Endogenous peroxidase activity was blocked with 0.3% H_2_O_2_/PBS. Slides were blocked for 4h with 10% goat serum. For the detection of neovessels, slides were incubated with 1° antibody for CD31 (ab28364, Abcam, 1:25 dilution) or VE-Cadherin (ab205336, Abcam, 1:100 dilution) in PBST overnight at 4°C, then incubated with HRP-conjugated goat anti-rabbit 2° antibody at 1:200 in PBST for 2h at room temperature. Positively stained neovessels were detected using a DAB Peroxidase Substrate Kit (Vector Labs), and counterstained with hematoxylin. For the detection of arterioles, slides were incubated with 1° antibody for α-SMA conjugated to alkaline phosphatase (A5691, Sigma-Aldrich, 1:100 dilution) in PBST overnight at 4°C. Positively stained arterioles were detected using Vector Red Alkaline Phosphatase Substrate Kit (Vector Labs) and counterstained with hematoxylin. Staining was quantified using ImageJ (NIH).

### Cell Culture and Treatments

Human coronary artery ECs (HCAECs; Cell Applications Inc.) were cultured in MesoEndo (Cell Applications Inc.) and used at passages 3 to 5. Cells were seeded at 1×10^5^ cells/well in 6-well plates and cultured for 24h at 37°C in 5% CO_2_ prior to any treatment. HCAECs were treated with rHDL, apoA-I, Glyc-rHDL, Ox-rHDL (20μmol/L, equivalent to 0.6mg/mL; final apoA-I concentration), PLPC vesicles (2mmol/L), or PBS control for 18h, then exposed to MesoEndo media with a baseline glucose concentration of 5.5mmol/L for normal glucose conditions, or supplemented with D-glucose to a final concentration 25mmol/L for high glucose conditions for 72h. Media was replenished every 24h. HCAECs were exposed to either normoxia or hypoxia (1.2% O_2_ balanced with N_2_) for 6h.

### Lentiviral shRNA SR-BI Knockdown in HCAECs

HCAECs were seeded at 5×10^4^ cells/well in 6-well microplates and cultured for 18h at 37°C. Cells were then transduced with 5×10^8^ lentiviral particles/mL containing shRNA for SR-BI (shSR-BI) for 24h. Media was replaced with fresh MesoEndo media and cells were allowed to grow for a further 4d. Five days post-transduction, cells were treated with rHDL (20μmol/L) or PBS for 18h then exposed to high glucose (25 mmol/L) for 72h, then hypoxia (1.2% O_2_) for 6h. SR-BI protein was determined by Western blotting by probing cell lysates with SR-BI antibody (ab52629, Abcam, 1:1000 dilution).

### siRNA-mediated PDK4 Knockdown in HCAECs

HCAECs were seeded at 1.5×10^5^ cells/well in 6-well microplates and cultured for 24h at 37°C. Media was replaced with 500µL of Opti-MEM, and 50nM siRNA for PDK4 or a scrambled control (Millenium Science Australia) was added in 300µL Opti-MEM containing 2.5% Lipofectamine 3000 (Thermo Fisher Scientific). Media was replaced after 6h, and cells were cultured for an additional 18h before further treatment or harvest.

### PDK4 Inhibition in Diabetic Mice

For these studies, we used PDK4 inhibitor ‘8c’ (PDK4-IN-1 hydrochloride, HY-135954A, MedChemExpress) with excellent specificity.^19^ The action of the PDK4 inhibitor on endothelial tubule formation was first assessed in vitro using the Matrigel tubulogenesis assay. Briefly, HCAECs were seeded at 1×10^5^ cells/mL in 6-well microplates and cultured for 24h at 37°C in 5% CO_2_. Cells were then treated with either PDK4 inhibitor (20μol/L) or control for 24h. Treated cells were then seeded at 1×10^4^ cells/mL on 50µL polymerized growth factor-reduced Matrigel (BD Biosciences) in 96-well microplates, then incubated at 37°C in normoxia for 6h.

In vivo, parallel cohorts of diabetic mice received topical applications of the chemical inhibitor, PDK4-IN-1 hydrochloride (50μg) in both wounds at the time of surgery then daily until day 5 post-wounding. For each mouse, one wound received topical rHDL (50μg/wound/day) or PBS (control) daily until final endpoint. Wound closure was assessed daily using microcalipers. Wound perfusion was tracked longitudinally by laser Doppler imaging at days 1, 3, 7 and 9.

### Cellular Metabolic Phenotype

The metabolic phenotype of pre-treated HCAECs was assessed using the Seahorse Bioanalyzer Cellular Metabolic Phenotype Assay (Agilent). After 48h glucose exposure, HCAECs were re-seeded across two 96-well microplates at 2×10^4^ cells/well in the corresponding glucose media. 24h later, media was changed to assay media and one microplate was run immediately (normoxia) whilst the second microplate was exposed to hypoxia for 6h. Immediately prior to running each plate, several empty wells were seeded with un-treated cells (2×10^4^ cells/well) to act as an interplate control. To account for the 6h difference in growth time, data from each well were normalized to total DNA content using the CyQUANT Cell Proliferation Assay (Thermo Fisher Scientific).

### Lactate Assay

Lactate was detected using the Colorimetric/Fluorometric L-Lactate Assay Kit (ab65330, Abcam). Cell media samples were diluted 1:20 in Assay Buffer, and combined with the enzyme and substrate reaction mix to generate colour, which was measured at 450nm using an iMark Microplate Absorbance Reader (Bio-Rad).

### Matrigel Tubulogenesis Assay

Pre-treated HCAECs were seeded at 1×10^5^ cells/mL on 50µL polymerized growth factor-reduced Matrigel (BD Biosciences) in 96-well microplates, then incubated at 37°C in normoxia or hypoxia for 6h. Wells were imaged at 5X magnification under light microscopy. Tubule and branch point number was determined using ImageJ (NIH).

### Boyden Chamber Migration Assay

Pre-treated HCAECs were seeded at 2×10^5^ cells/mL in Boyden Chamber transwells in Opti-MEM (Thermo Fisher Scientific) containing recombinant VEGFA protein (10ng/mL), then incubated for 18h. Membranes were excised, stained with DAPI and migrated cells imaged at 10X magnification using fluorescence microscopy. Cell number was determined using ImageJ (NIH).

### RNA Expression

Tissues were homogenised at 6000 x g in 500µL of TRI reagent using a Precellys homogeniser. For treated HCAECs, cells were washed with ice-cold PBS and scraped in 250µL TRI reagent. Following cell lysis, total RNA was isolated as previously described.^20^ 500ng of RNA was reverse transcribed using the iScript cDNA Synthesis Kit (BioRad). For quantitative real-time PCR, reactions were performed in duplicate with primers used at 10μM in combination with the iQ SYBR green Supermix (Bio-Rad) in a Bio-Rad thermocycler. The protocol conditions included: initial denaturation and enzyme activation at 95°C for 3 minutes; 40 cycles of denaturation at 95°C for 30 seconds, annealing at 60°C for 30 seconds, followed by extension at 72°C for 30 seconds. A non-template control (NTC) was included for each gene. qPCR was performed for human or murine *PDK4/Pdk4*, human β2-microglobulin (B2M), and murine *36B4* (Supplementary Table 1 for primer sequences) in HCAECs or wound tissue. Relative changes in gene expression were normalized using the ^ΔΔ^Ct method to human *B2M* or murine *36B4*.

### Protein Expression

Tissues were homogenised at 6000 x g in 150µL of radioimmunoprecipitation (RIPA) buffer for western blot analysis or Cell Extraction Buffer for ELISA analysis using a Precellys homogeniser. For treated HCAECs, cells were washed with ice-cold PBS and scraped in 100µL RIPA buffer. Protein concentration was determined using the Pierce BCA Protein Assay Kit (Life Technologies). Whole-cell protein extracts were probed with antibodies for various targets. Even protein loading was confirmed by probing for α-tubulin (Supplementary Table 2 for antibody details). Human and murine PDK4 expression was measured by ELISA (ab126582 and ab215544, Abcam), as was human HIF-1α (RDSDYC19352, R&D Systems).

### Chromatin Immunoprecipitation Assay

Following cell treatments performed in 6-well microplates, proteins and DNA were crosslinked by addition of 37% formaldehyde (final concentration of 1%). After 10min, 1M ice-cold glycine was added. Cells were washed and scraped in ChIP lysis buffer. DNA was sonicated to 300bp fragments. Chromatin samples were pre-cleared by incubation with ChIP-grade protein G magnetic beads for 2h at 4°C. Samples were incubated with an antibody for FOXO1 or an IgG Normal Rabbit control (Supplementary Table 2 for antibody details) overnight at 4°C. Protein was immunoprecipitated using the protein G magnetic beads, washed 3x with low salt buffer and 1x with high salt buffer before elution of DNA. Crosslinks were reversed with 5M NaCl, and Proteinase K. DNA was purified using a QIAquick PCR clean-up kit (Qiagen), and RT-qPCR performed to amplify the insulin-response sequence 1 (IRS1) in the PDK4 promoter region using the same method as detailed in the RNA Expression section (See Supplementary Table 1 for primer sequences). Expression data was compared to a 1% chromatin input control and normalised to the IgG-only control to calculate the fold enrichment of the target sequence.

### Statistical Analysis

Data are expressed as mean±SEM. Within experiments, each experimental condition was performed in triplicate. Each whole experiment was then completed at least three times. A two-way ANOVA was used when comparing data across multiple time points or when comparing data spanning three or more conditions (Tukeys posthoc test of significance). Differences between treatment groups were calculated using a Student’s t-test or one-way ANOVA (Tukeys posthoc test of significance). Paired t-tests were used when comparing data points from the same mice. Significance was set at *P*<0.05.

## RESULTS

### Diabetes impairs metabolic reprogramming responses to wound ischemia and is rescued by rHDL in diabetic mice

Mice receiving STZ injections had significantly higher blood glucose levels than non-diabetic control mice (Non-diabetic: 10.1±1.3 versus Diabetic: 25.7±6.4 mmol/L, *P*<0.0001) and lower body weights (Non-diabetic: 24.5±1.8 versus Diabetic: 22.9±2.1 g, *P*<0.05) at the time of wound surgery (Supplementary Table 3). At day 1 post-wounding in non-diabetic mice there was an induction of PDK4 protein levels (3.0±1.2 to 9.3±2.0 ng/mL, 207% increase, *P*<0.05, Fig. 1A). This induction did not occur in diabetic mice. However, topical rHDL restored the induction of PDK4 protein expression at day 1 post-wounding in diabetic wounds (4.4±1.0 to 7.4±1.6 ng/mL, 68% increase, *P*<0.05, Fig. 1A). Consistent with this, at day 3 post-wounding rHDL-treated wounds exhibit enhanced pPDC in diabetic mice (63±8 to 167±39 %, 165% increase, *P*<0.01, Fig. 1B-C). In non-diabetic mice, wound *Pdk4* mRNA levels were also significantly increased at day 1 post-wounding (100±43 to 288±75 %, 188% increase, *P*<0.05, Fig. 1D). Diabetes impaired this induction, with *Pdk4* mRNA expression significantly reduced in Day 1 diabetic wounds (288±75 to 127± 29 %, 56% decrease, *P*<0.01, Fig. 1D). In diabetic mice, topical application of rHDL enhanced *Pdk4* mRNA levels at day 3 post-wounding (22±5 to 40±9 %, 82% increase, *P*<0.05, Fig. 1D and inset). When tested in female mice, topical rHDL caused a non-significant increase in *Pdk4* mRNA levels in diabetic wounds at day 1 post-wounding, compared to day 1 diabetic PBS wounds (Diabetic rHDL: 135.2±21.9% vs. Diabetic PBS: 82.7±17.2%, P=0.074) (Supplementary Fig. 2A, Supplementary Table 4).

**Figure 1.**
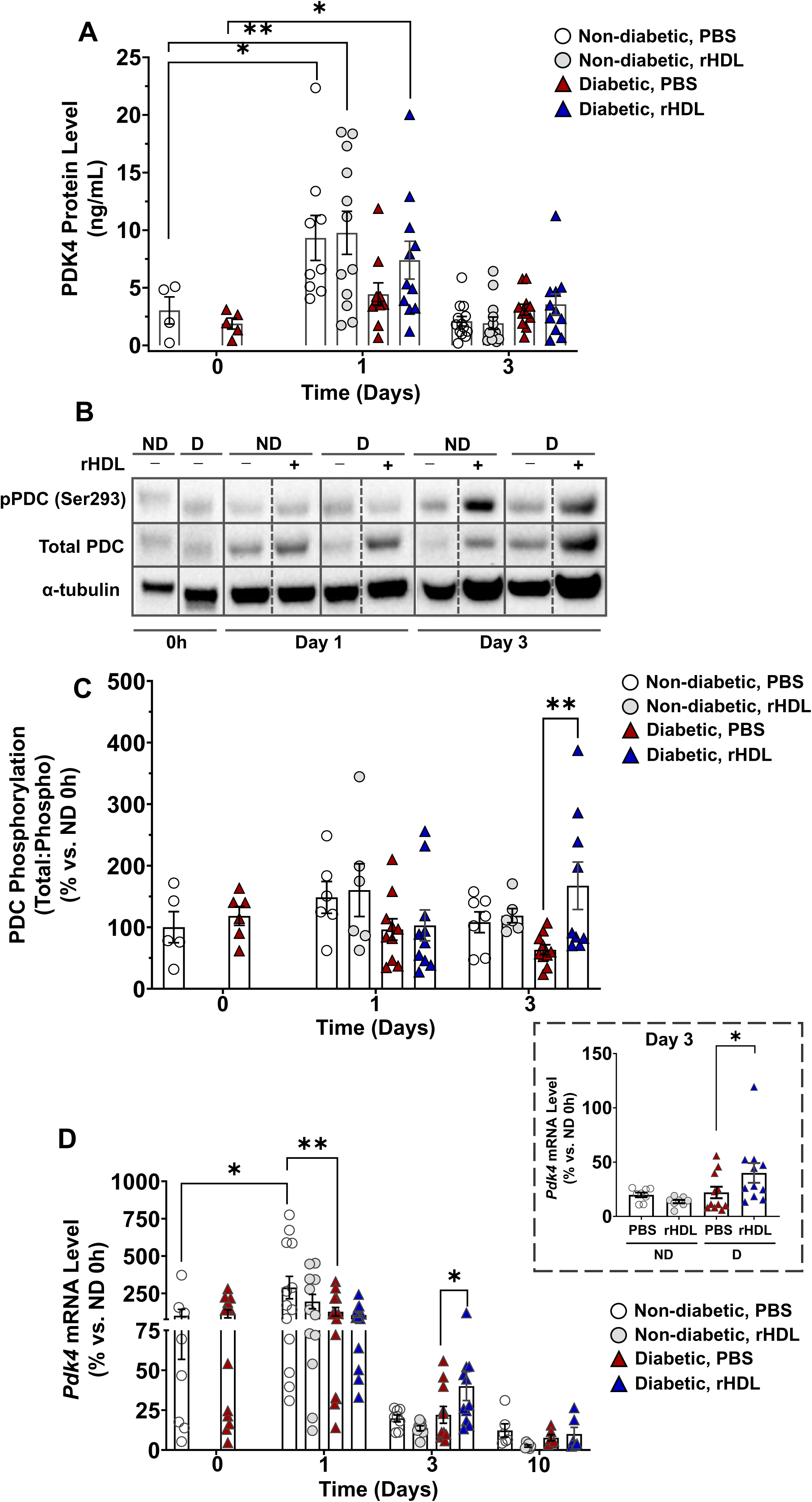
Diabetes impairs metabolic reprogramming response to wound ischemia and rHDL rescues this impairment in diabetic mice. Two full-thickness wounds were created on non-diabetic and diabetic C57Bl/6J mice. Mice received daily topical applications of rHDL (50 μg/wound) or PBS (vehicle). Wound tissue was collected from non-diabetic (white and grey circles) and diabetic (red and blue triangles) mice at days 0, 1, 3, and 10 post-wound creation. (A) PDK4 protein expression at days 0, 1 and 3 post-wounding was measured in wound lysates by ELISA (n=4-12) \**P*<0.05, \*\**P*<0.01 vs controls by two-way ANOVA. (B-C) Phosphorylation of the PDC at days 0, 1, and 3 post-wounding was measured in wounds by western blotting (n=5-10) \*\**P*<0.01 vs. PBS control by two-way ANOVA. (D) *Pdk4* mRNA expression was measured in wounds using qRT-PCR. Gene expression was normalized using the ^ΔΔ^Ct method to murine *36B4.* (n=8-13) \**P*<0.05, \*\**P*<0.01 vs. control by two-way ANOVA. \**P*<0.05 by paired t-test. Data are expressed as mean±SEM.

### rHDL increases wound neovascularization and rescues diabetes-impaired healing *in vivo*

Blood flow perfusion of the wound was determined as a ratio of rHDL:PBS corrected against day 1 using laser Doppler imaging. In diabetic mice, rHDL significantly increased wound blood flow perfusion 3 days post-wounding (100±2 to 132±24 %, 32% increase, *P*<0.05, Fig. 2A). At Day 10 post-wounding, the PBS-treated wounds of diabetic mice contained fewer CD31^+^ neovessels compared to the PBS-treated wounds of non-diabetic mice (36±4 to 21±3 neovessels, 42% decrease, *P*<0.05, Fig. 2B). rHDL treatment rescued this impairment in diabetic mice (21±3 to 34±4 neovessels, 62% increase, *P*<0.05). Diabetic PBS wounds also had fewer VE-Cadherin^+^ neovessels compared to non-diabetic PBS wounds (291±27 to 173±17 neovessels, 42% decrease, *P*<0.05, Supplementary Fig. 3A). However, no significant differences were observed between diabetic rHDL and non-diabetic PBS wounds. No differences were found in α-SMA^+^ arteriole numbers across all groups (Supplementary Fig. 3B). The PBS-treated wounds of diabetic mice exhibited reduced rates of wound closure, reaching significance at Day 8 (60±2 to 48±4 % wound closure, 20% decrease, *P*<0.05, Fig. 2E-F), compared to non-diabetic mice with PBS-treated wounds. Topically applied rHDL significantly increased the rate of wound closure in diabetic mice across all timepoints (increases ranging from 31.9-154.9%, *P*<0.01, *P*<0.0001), compared to diabetic PBS-treated wounds.

**Figure 2.**
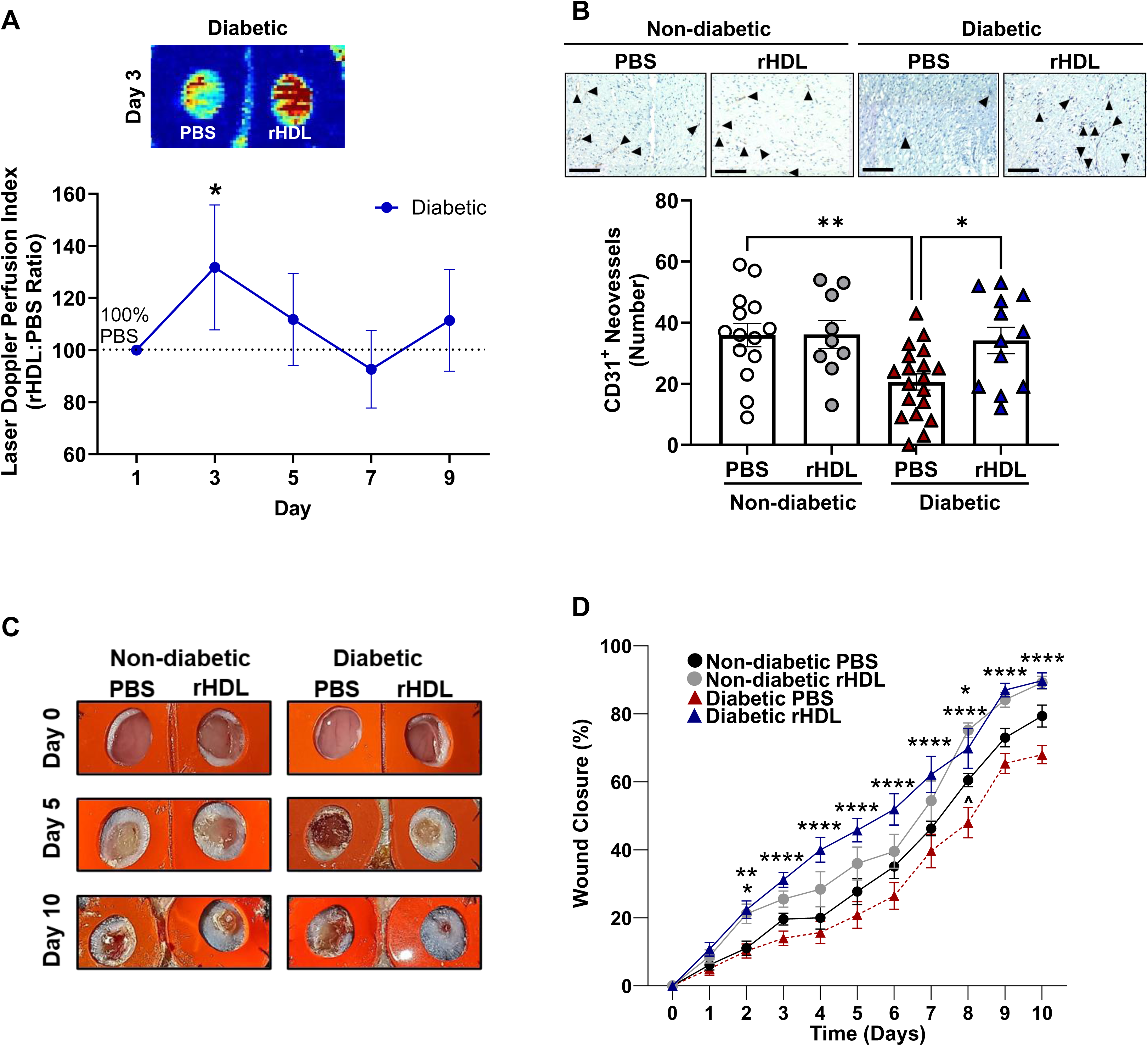
rHDL increases wound neovascularization and rescues diabetes-impaired wound healing in vivo. Two full-thickness wounds were created on non-diabetic and diabetic C57Bl/6J mice. Mice received daily topical applications of rHDL (50 µg/wound) or PBS (vehicle). (A) The rHDL:PBS wound blood flow perfusion ratio was determined using laser Doppler perfusion imaging; images represent high (red) to low (blue) blood flow from day 1-9 in non-diabetic (black circles) and diabetic (blue circles) mice. \**P*<0.05 vs. PBS control by paired t-test. Neovessels were identified in wound sections using immunohistochemistry for (B) CD31. Photomicrographs represent wounds stained for CD31 (stained brown, denoted by arrows). Scale bars, 200 μm. \**P*<0.05, \*\**P*<0.01 by one-way ANOVA (n=9-14). (C-D) Wound area was calculated from the average of three daily diameter measurements along the x-, y-, and z-axes. Wound closure is expressed as a percentage of initial wound area at day 0. \**P*<0.05, \*\**P*<0.01, \*\*\**P*<0.001, \*\*\*\*\**P*<0.0001 vs. relevant controls by two-way ANOVA. ^*P*<0.05 vs. non-diabetic PBS-treated wound by two-way ANOVA. (n=12). Non-diabetic PBS-treated wounds (black circles), non-diabetic rHDL-treated wounds (grey circles), diabetic PBS-treated wounds (red triangles), diabetic rHDL-treated wounds (blue triangles). Data are expressed as mean±SEM.

In female mice, laser Doppler analyses revealed that the rHDL:PBS ratio was significantly increased at both days 1 and 3 post-wounding, with the highest levels observed at day 1 (Supplementary Fig. 2B-C).

### High glucose impairs metabolic reprogramming responses to hypoxia, which is restored by rHDL *in vitro*

We next studied how the PDK4/PDC axis was affected in high glucose and hypoxia *in vitro*, and its regulation by rHDL. Under normal glucose conditions at 5mM, hypoxia exposure increased EC *PDK4* mRNA expression (100±5 to 165±16 %, 65% increase, *P*<0.05, Fig. 3A). However, exposure to high glucose at 25mM did not elicit a further stepwise induction of *PDK4* in response to hypoxia. An additional stepwise induction is essential to adequately reduce the activity of the PDC to protect against both the effects of hypoxia *and* high glucose. However, preincubation with rHDL in high glucose restored this response to hypoxia, inducing *PDK4* mRNA (176±15% to 240±17%, 37% increase, *P*<0.05), compared to the PBS high glucose control. Assessment of the effects of individual components of rHDL, i.e. lipid-free apoA-I and 1-palmitoyl-2-linoleoyl-phosphatidylcholine (PLPC) showed that *PDK4* expression in PLPC-treated cells was significantly higher than the PBS control cells (PLPC: 128±9% vs. PBS: 100±3%, *P*<0.05, Fig. 3B). No differences were observed between apoA-I treated cells and PBS control cells. We also evaluated the effects of rHDL modified by glycation or oxidation. *PDK4* mRNA levels were elevated in cells treated with either glycated (143±8%, *P*<0.001) or oxidized (130±7%, *P*<0.05) rHDL. Consistent with the mRNA findings, rHDL increased PDK4 protein (7.4±0.6 to 10.8±0.8 ng/mL, 44% increase, *P*<0.05, Fig. 3C), compared to the PBS high glucose control. Furthermore, in normal glucose conditions, PDC phosphorylation (pPDC) increased in response to hypoxia (130±11 to 213±25 %, 64% increase, *P*<0.01, Fig. 3D-E). Exposure to high glucose in hypoxia did not induce a further elevation in pPDC levels, however, rHDL corrected this, and promoted higher pPDC levels (100±6 to 170±22 %, 70% increase, *P*<0.01), compared to PBS treated ECs in high glucose and hypoxia.

**Figure 3.**
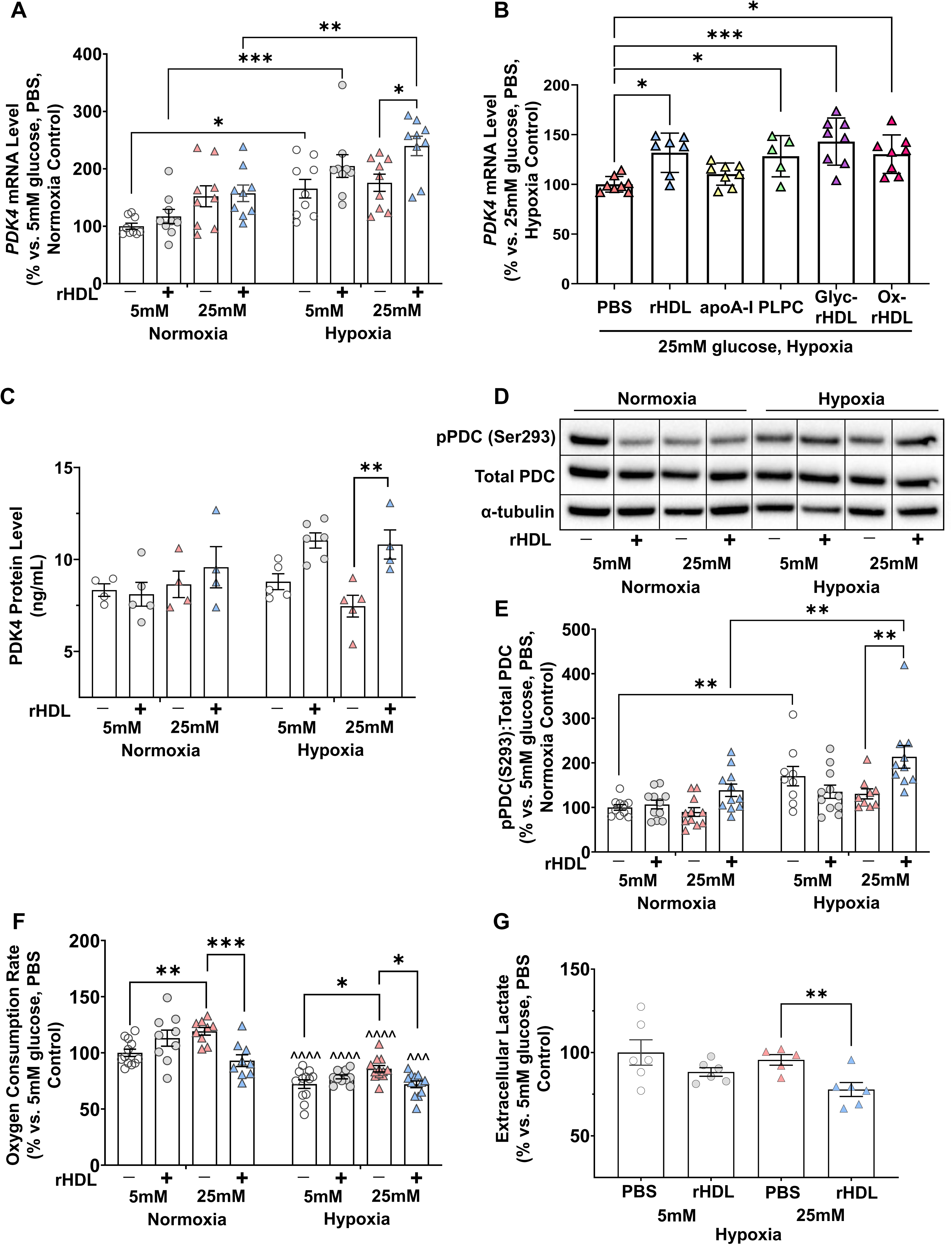
High glucose impairs metabolic reprogramming responses to hypoxia in ECs and rHDL restores this impairment *in vitro*. HCAECs were incubated with rHDL (20 µM) or PBS (vehicle control) for 18 h, then exposed to normal (5 mmol/L,) or high (25 mmol/L) glucose conditions for 72 h. Following this, cells were exposed to normoxia or hypoxia (1.2% O_2_) for 6 h. (A) *PDK4* mRNA expression was measured using qRT-PCR and normalized using the ^ΔΔ^Ct method to human *B2M,* (n=9). (B) *PDK4* mRNA expression was measured in HCAECs incubated with rHDL, apoA-I, glycated rHDL (Glyc-rHDL), oxidized rHDL, PLPC or PBS for 18 h then exposed to high glucose and hypoxic conditions (n=6-8). (C) PDK4 protein expression was measured using an ELISA, (n=4-6). (D-E) Phosphorylation of the PDC at serine 293 was measured using western blotting, with data expressed as the ratio of pPDC(S293):Total PDC, and then normalized to α-tubulin, (n=10). \**P*<0.05, \*\**P*<0.01, \*\*\**P*<0.001 by two-way ANOVA. (F) Oxygen consumption of treated cells was measured using the Seahorse Bioanalyser system, with data from each well normalized to cell number (n=9-12). \**P*<0.05, \*\**P*<0.01, \*\*\**P*<0.001 by two-way ANOVA. ^^^*P*<0.001, ^^^^*P*<0.0001 vs. normoxia control by two-way ANOVA. (G) Lactate in the media of hypoxia-exposed and rHDL treated cells was measured using a colorimetric assay, (n=5, 6). \*\**P*<0.01 by unpaired t-test. 5 mM glucose, PBS (white circles), 5 mM glucose, rHDL (grey circles), 25 mM glucose, PBS (red triangles), 25 mM glucose, rHDL (blue triangles). Data are expressed as mean±SEM.

We next aimed to determine whether this change in regulation of the PDK4/PDC axis was associated with changes in cellular oxygen consumption rate (OCR), a measure of mitochondrial respiration ^21^. In normoxia, high glucose exposure increased OCR (100±3 to 119±3 %, 19% increase, *P*<0.01, Fig. 3F), compared to normal glucose. However, rHDL prevented this and kept OCR at baseline levels. Overall, hypoxia lowered the OCR across all treatments when compared to normoxia groups. However, high glucose exposure in hypoxia increased OCR (72±4 to 86±3 %, 19% increase, *P*<0.05), compared to the normal glucose control. rHDL prevented this high glucose-induced elevation of the OCR in hypoxia, significantly lowering the OCR (86±3 to 72±3 %, 16% decrease, *P*<0.05), compared to PBS control cells in high glucose and hypoxia. This suggests rHDL restores adequate suppression of mitochondrial respiration in response hypoxia when disrupted by high glucose. In hypoxia, rHDL treatment also significantly decreased extracellular lactate under high glucose conditions (96±3 to 78±4 %, 19% decrease, *P*<0.01, Fig. 3G), compared to the PBS control, further supporting that rHDL suppresses metabolic activity in hypoxia and high glucose.

### rHDL rescues high glucose-impaired EC function *in vitro*

In hypoxia, high glucose exposure impaired tubule (117±4 to 85±9 %, 28% decrease, Fig. 4A-B) and branch point formation (128±2 to 95±7 %, 26% decrease, Fig. 4A,C) (*P*<0.05 for both), compared to cells in normal glucose and hypoxia. rHDL rescued this impairment, increasing tubule number (85±7 to 136±10 %, 60% increase, *P*<0.001) and branch points (95±7 to 169±4 %, 78% increase, *P*<0.0001). High glucose impaired migration in normoxia (100±6 to 60±7, 40% decrease, *P*<0.01, Fig. 4D-E) and hypoxia (107±7 to 65±10, 39% decrease, *P*<0.01), compared to the respective controls. In hypoxia, rHDL rescued migration, elevating the number of migrated cells (65±10 to 100±6 %, 55% increase, *P*<0.05), compared to the PBS control.

**Figure 4.**
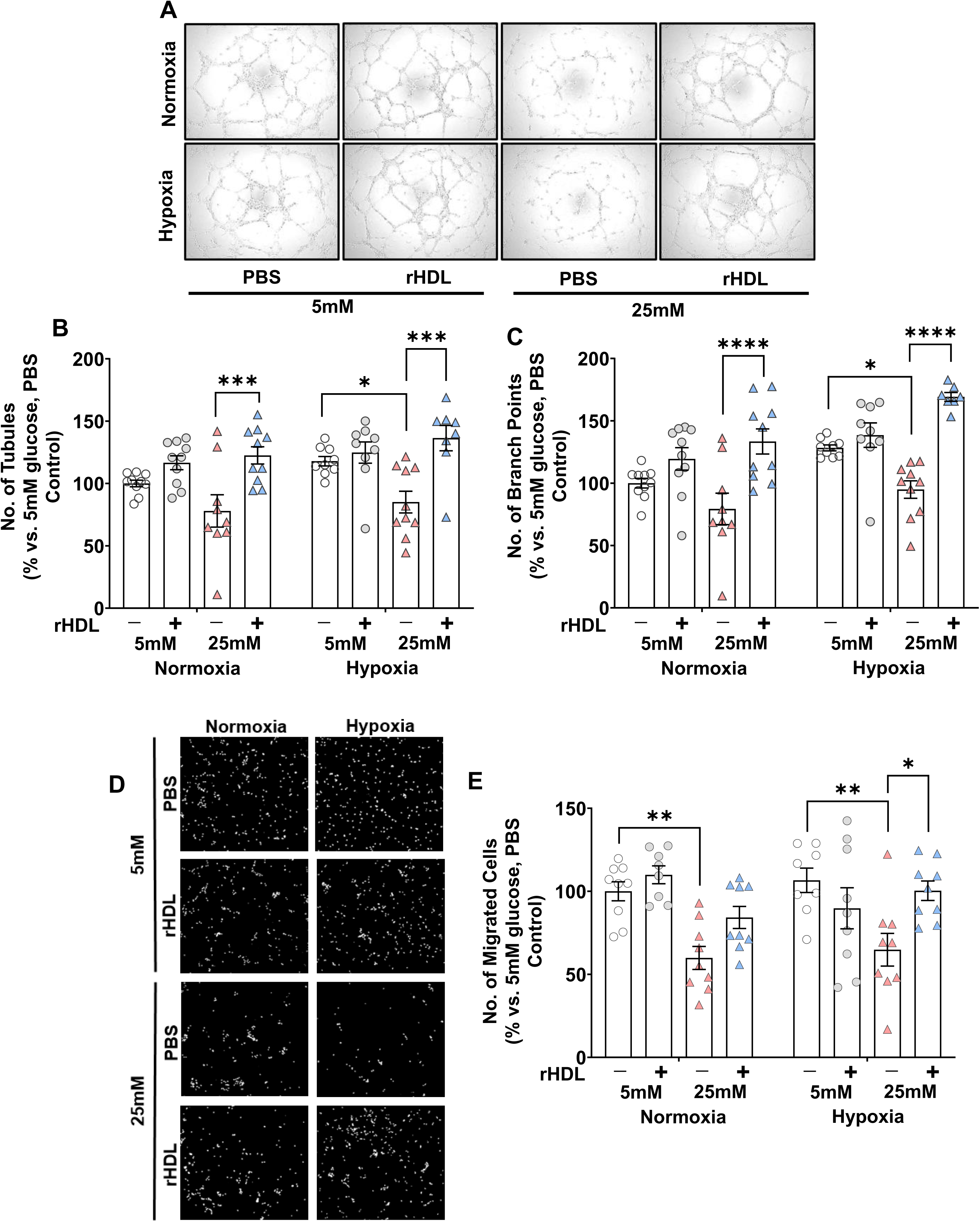
rHDL rescues high glucose-impaired EC function *in vitro*. HCAECs were incubated with rHDL (20 µM) or PBS (vehicle control) for 18 h, then exposed to normal (5 mmol/L) or high (25 mmol/L) glucose conditions for 72 h. (A) Treated cells were then seeded on Matrigel and exposed to normoxia or hypoxia (1.2% O_2_) for 6 h. Representative images of tubules from all conditions. Matrigel tubules (B) and branch points (C) were imaged using light microscopy and quantified using ImageJ software (n=9). Treated cells were seeded in wells containing a permeable membrane to measure cell migration. (D) Representative images from all conditions of imaged membranes of migrated DAPI-stained cells by fluorescence microscopy and quantified (E) using ImageJ software (n=9). \**P*<0.05, \*\**P*<0.01, \*\*\**P*<0.001, \*\*\*\**P*<0.0001 by two-way ANOVA. 5 mM glucose, PBS (white circles), 5 mM glucose, rHDL (grey circles), 25 mM glucose, PBS (red triangles), 25 mM glucose, rHDL (blue triangles). Data are expressed as mean±SEM.

### SR-BI knockdown attenuates the induction of *PDK4* by rHDL

We next used a lentiviral shRNA knockdown approach to determine the importance of SR-BI in mediating the effects of rHDL on *PDK4*. SR-BI protein expression was significantly lower in SR-BI cells (42±3%) compared to Control cells (100±3, *P*<0.0001, Fig. 5A). In Control cells exposed to high glucose and hypoxic conditions, rHDL augmented *PDK4* mRNA levels compared to the PBS cells (Control rHDL: 130±6% versus Control PBS: 100±3%, *P*<0.001, Fig. 5B). However, this induction was abrogated in SR-BI cells.

**Figure 5.**
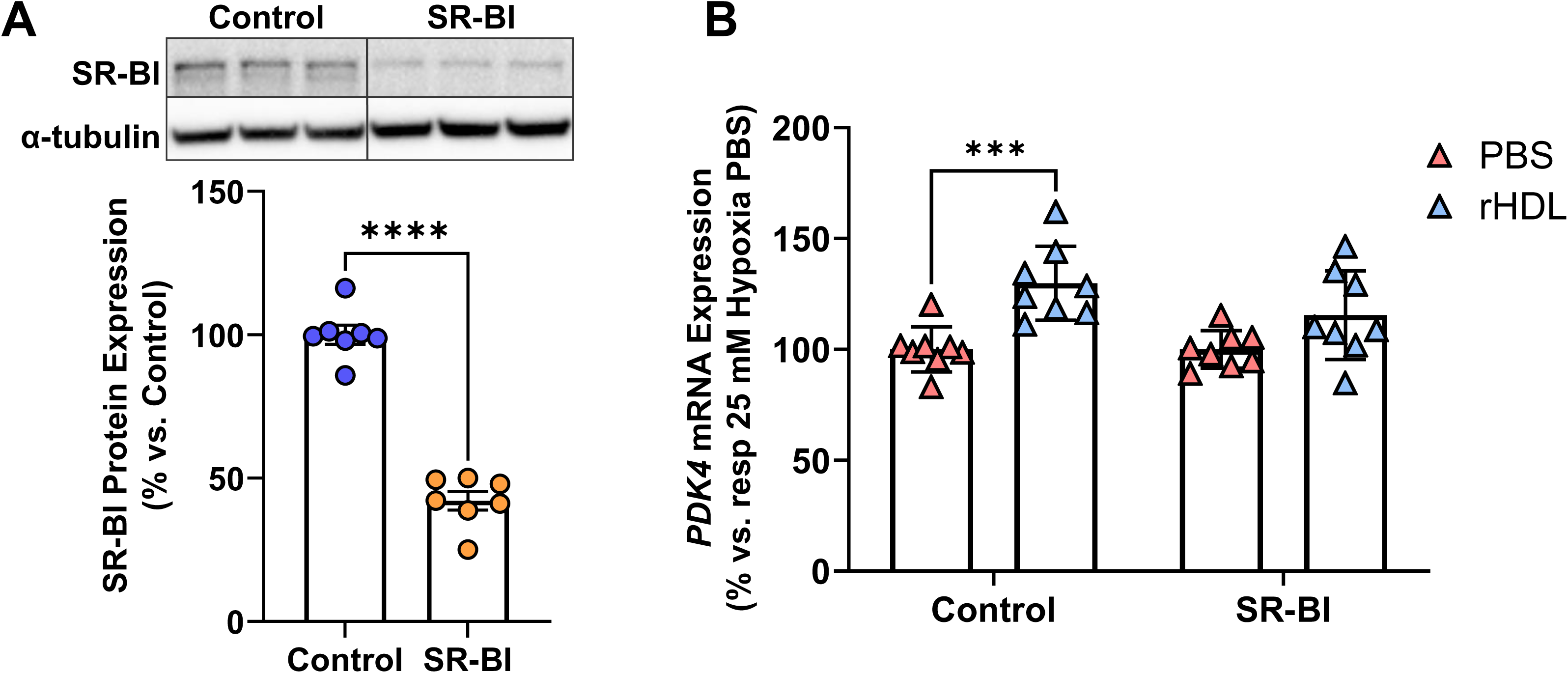
SR-BI knockdown attenuates the induction of *PDK4* by rHDL. HCAECs were transduced with lentiviral particles containing shRNA for SR-BI (SR-BI) for 24h, then incubated with fresh media and culture for a further 4d. Five days post-transduction, cells were treated with rHDL (20μmol/L) or PBS for 18h then exposed to high glucose (25 mmol/L) for 72h, then hypoxia (1.2% O_2_) for 6h. (A) SR-BI protein was measured using western blotting densitometry, with data normalized to α-tubulin. \*\*\*\**P*<0.0001 by unpaired t-test. (B) *PDK4* mRNA expression was measured using qRT-PCR and normalized using the ^ΔΔ^Ct method to human *B2M,* (n=8). Data are expressed as mean±SEM.

### PDK4 knockdown attenuates the pro-angiogenic effects of rHDL

High glucose exposure impaired tubule formation in the non-transfected (100±3 to 81±6 %, 19% decrease, *P*<0.05, Fig. 6B), scrambled siRNA control (97±3 to 71±4 %, 27% decrease, *P*<0.01) and PDK4 siRNA ECs (72±7 to 63±6 %, 12% decrease, *P*<0.05), when compared to their respective normal glucose controls (Fig. 6A-B). PDK4 knockdown impaired tubule formation in normal glucose PBS cells, compared to both the non-transfected (100±3 to 72±7 %, 28% decrease, *P*<0.001) and scrambled siRNA controls (97±3 to 72±7 %, 26% decrease, *P*<0.01). rHDL treatment rescued high glucose-impaired tubulogenesis in both non-transfected (81±6 to 121±3 %, 50% increase, *P*<0.0001) and scrambled siRNA controls (71±4 to 108±6 %, 52% increase, *P*<0.0001). However, in the cells deficient in PDK4, the proangiogenic effects of rHDL on tubule number in high glucose were completely attenuated.

**Figure 6.**
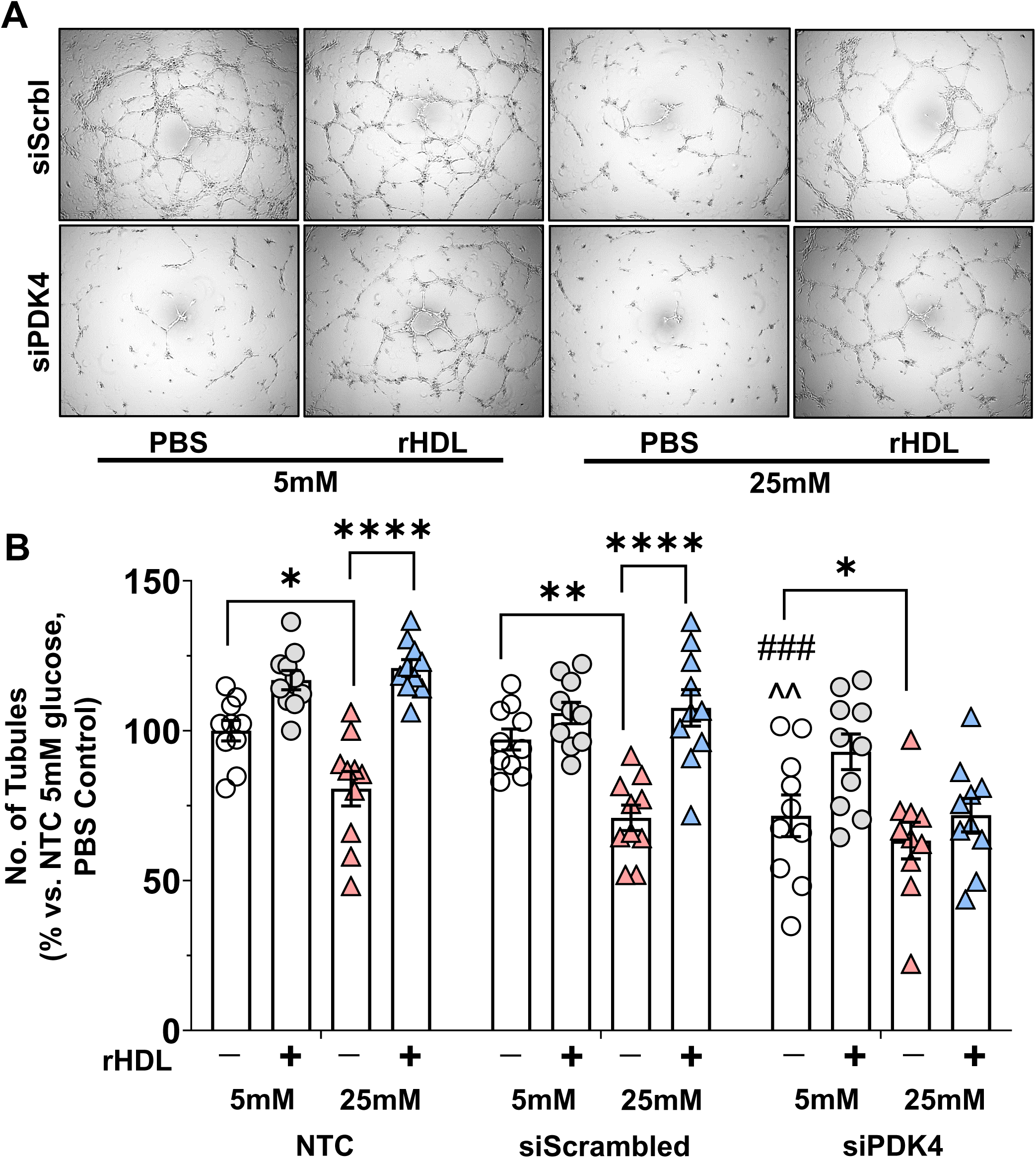
PDK4 knockdown attenuates the pro-angiogenic effects of rHDL *in vitro*. HCAECs were treated with PDK4-specific siRNA (50 nM) or a scrambled control for 6 h, then incubated with fresh media for 48 h. HCAECs were then incubated with rHDL (20 µM) or PBS (vehicle control) for 18 h, then exposed to normal (5 mmol/L) or high (25 mmol/L) glucose conditions for 72 h. Treated cells were then seeded on Matrigel and exposed to normoxia or hypoxia (1.2% O_2_) for 6 h. (A) Representative light microscopy images of tubules from the siScrambled and siPDK4 conditions. (B) Matrigel tubules were imaged and quantified using ImageJ software (n=10). \**P*<0.05, \*\**P*<0.01, \*\*\*\**P*<0.0001 vs. controls by two-way ANOVA. ^###^*P*<0.001 vs. non-transfected control by two-way ANOVA. ^^^^*P*<0.01 vs. siScrambled control by two-way ANOVA. 5 mM glucose, PBS (white circles), 5 mM glucose, rHDL (grey circles), 25 mM glucose, PBS (red triangles), 25 mM glucose, rHDL (blue triangles). Data are expressed as mean±SEM.

We next determined if these effects occurred *in vivo* in diabetic mice. For these studies, we used a PDK4 inhibitor (PDK4i) with excellent specificity as previously described and characterized.^19^ We accordingly confirmed that treatment with PDK4i in endothelial cells significantly inhibited *in vitro* angiogenesis (Supplementary Figure 4). *In vivo*, in diabetic mice, rHDL treatment once again promoted wound closure (Fig. 7A) and wound perfusion early at day 3 post-wounding when angiogenesis is important (Fig. 7B). However, daily application of the PDK4 inhibitor in the first 5 days post-wounding attenuated the ability of rHDL to rescue diabetes-impaired wound closure and wound perfusion.

**Figure 7.**
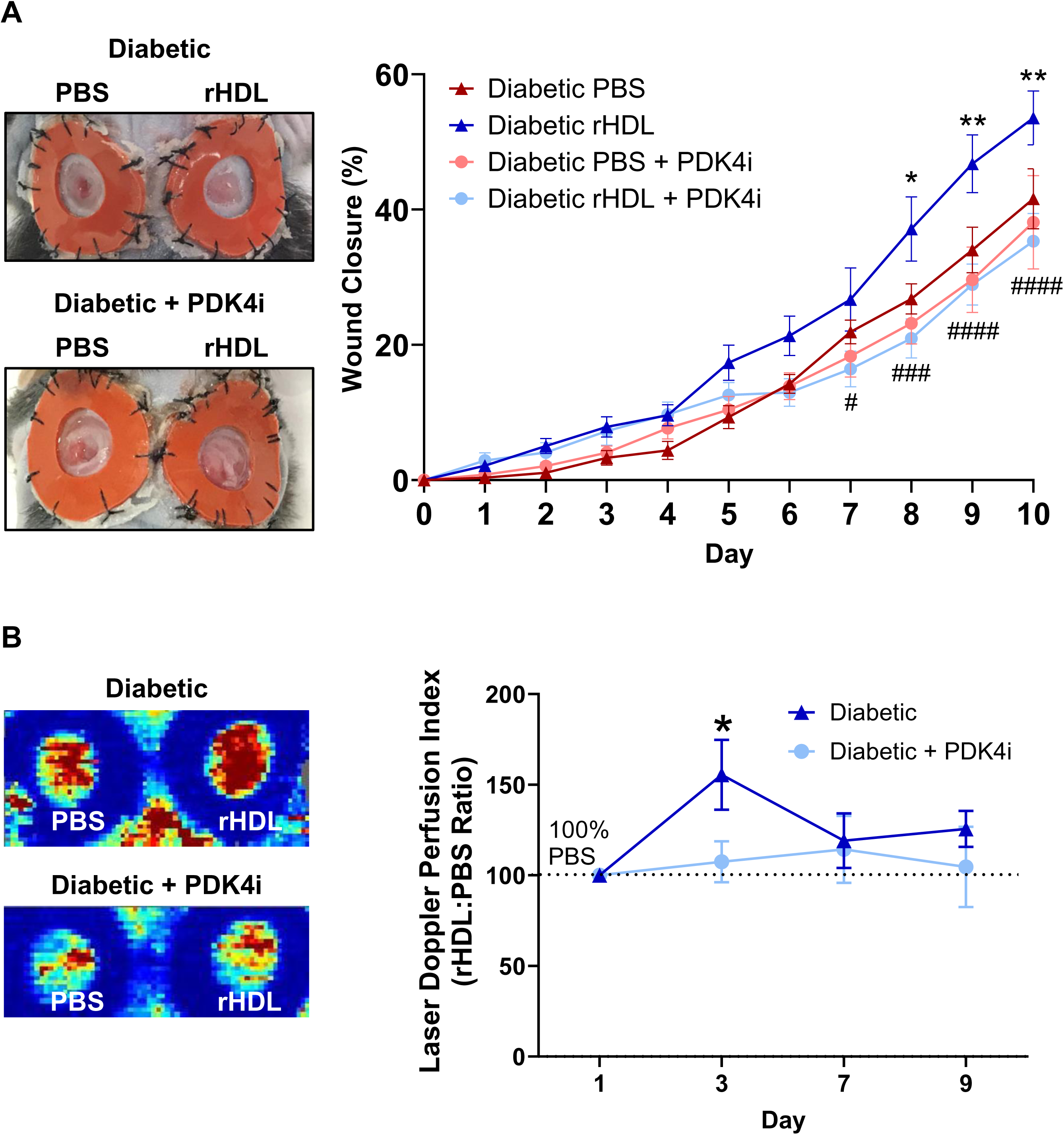
PDK4 inhibition attenuates the pro-angiogenic effects of rHDL *in vivo*. Two full-thickness wounds were created on diabetic C57Bl/6J mice. Mice received topical applications of the chemical inhibitor, PDK4-IN-1 hydrochloride (50μg) in both wounds at the time of surgery then daily until day 5 post-wounding. For each mouse, one wound received topical rHDL (50μg/wound/day) or PBS (control) daily until final endpoint. (A) Wound area was calculated from the average of three daily diameter measurements along the x-, y-, and z-axes. Wound closure is expressed as a percentage of initial wound area at day 0. \**P*<0.05, \*\**P*<0.01, diabetic rHDL vs. diabetic PBS; ^#^*P*<0.05, ^###^*P*<0.001, ^####^*P*<0.0001, diabetic rHDL + PDK4i vs. diabetic rHDL by two-way ANOVA. (n=8-9). Diabetic PBS-treated wounds (red triangles), diabetic rHDL-treated wounds (blue triangles), diabetic PBS + PDK4i-treated wounds (pink circles), diabetic rHDL + PDK4i-treated wounds (light blue circles). Data are expressed as mean±SEM. (B) The rHDL:PBS wound blood flow perfusion ratio was determined using laser Doppler perfusion imaging; images represent high (red) to low (blue) blood flow from day 1-9 in diabetic (blue triangles) and diabetic + PDK4i (light blue circles) mice. \**P*<0.05 vs. PBS control by paired t-test.

### High glucose suppresses the induction of HIF-1α in response to hypoxia, yet rHDL has no effect on HIF-1α but reduces PHD2 and PHD3 in hypoxia and high glucose

HIF-1α has been demonstrated to play a role in regulation of PDK4 transcription.^12^ Expression of HIF-1α was significantly increased with hypoxia exposure across all conditions as anticipated (*P*<0.0001, Fig. 8A). High glucose exposure impaired expression of HIF-1α under hypoxic conditions (83±6 to 62±6 pg/mL, 29% decrease, *P*<0.01), compared to normal glucose in hypoxia. rHDL treatment had no effect on HIF-1α expression in either normal or high glucose. The role of the PHD proteins (1, 2 and 3) is primarily to target HIF-1α for degradation in normoxia, with their expression downregulated in hypoxia to allow accumulation of HIF-1α.^22^ The PHD proteins also play roles in other transcription pathways.^10,23,24^ Despite rHDL causing no change in HIF-1α, rHDL caused a decrease in PHD1 in normoxia and high glucose (112±12 to 83±6 %, 26% decrease, *P*<0.05, Fig. 8B) but no effect in hypoxia (Fig. 8C). Whilst rHDL elicited no changes in PHD2 in normoxia (Fig. 8D), rHDL decreased PHD2 protein levels in high glucose conditions and in hypoxia, compared to PBS high glucose controls (99±10 to 73±6 %, 26% decrease, *P*<0.05, Fig. 8E). In normoxia, PHD3 expression was decreased with high glucose (100±4 to 80±7 %, 20% decrease, *P*<0.05, Fig. 8F). rHDL treatment further reduced PHD3 expression in high glucose in both normoxia (80±7 to 54±5 %, 32% decrease, *P*<0.01, Fig. 8G) and hypoxia (65±6 to 46±5 %, 29% decrease, *P*<0.05), compared to PBS high glucose-treated cells.

**Figure 8.**
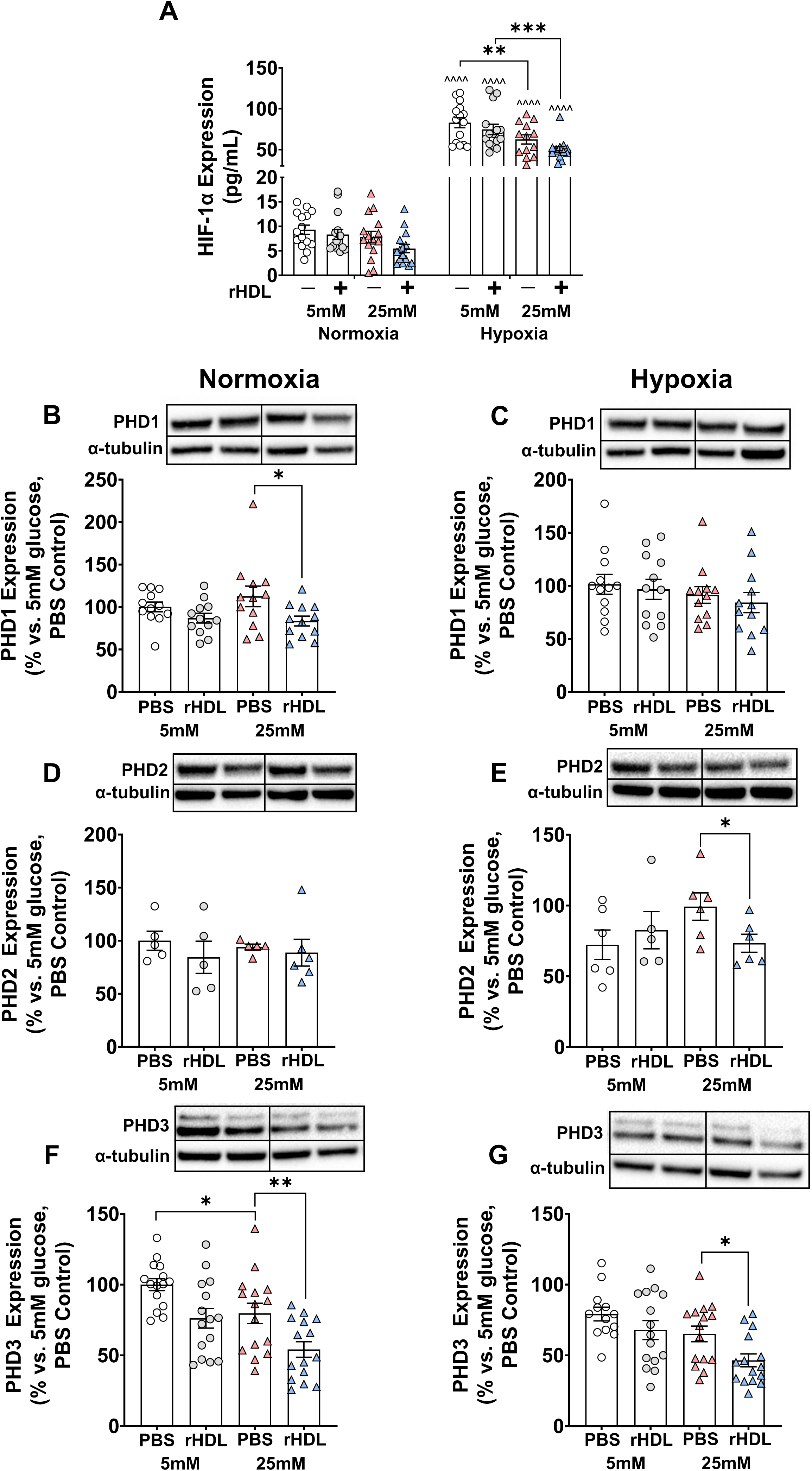
High glucose suppresses the induction of HIF-1α in response to hypoxia, rHDL has no effect on HIF-1α, but reduces expression of PHD1, PHD2, and PHD3. HCAECs were incubated with rHDL (20 µM) or PBS (vehicle control) for 18 h, then exposed to normal (5 mmol/L) or high (25 mmol/L) glucose conditions for 72 h. Following this, cells were exposed to normoxia or hypoxia (1.2% O_2_) for 6 h. (A) Whole-cell HIF-1α protein expression was measured using an ELISA (n=14). \*\**P*<0.01, \*\*\**P*<0.001 vs. controls. ^^^^*P*<0.0001 vs. normoxia controls, by two-way ANOVA. Whole-cell protein expression of (B-C) PHD1 (n=12), (D-E) PHD2 (n=5), and (F-G) PHD3 (n=14) was measured using western blotting densitometry, with data normalized to α-tubulin. \**P*<0.05, \*\**P*<0.01 by unpaired t-test. 5 mM glucose, PBS (white circles), 5 mM glucose, rHDL (grey circles), 25 mM glucose, PBS (red triangles), 25 mM glucose, rHDL (blue triangles). Data are expressed as mean±SEM.

### rHDL reduces phosphorylation of FOXO1 and enhances binding to the PDK4 promoter in high glucose and hypoxia

In the absence of a change in HIF-1α with rHDL, we next examined FOXO1, a transcription factor which interacts with the PDK4 promoter^25^ and plays a role in the regulation of angiogenesis.^26^ Phosphorylation of FOXO1 causes its nuclear exclusion and reduces its transcriptional activity. In hypoxia under normal glucose conditions, FOXO1 phosphorylation was increased (100±4 to 149±18 %, 49% increase, *P*<0.05, Fig. 9A-B), compared to the normoxia control. In hypoxia under high glucose, rHDL treatment decreased the phosphorylation of FOXO1 (193±13 to 138±15 %, 29% decrease, *P*<0.05), compared to PBS and high glucose-treated cells, returning it to the level observed under normal glucose conditions. A chromatin immunoprecipitation assay (ChIP) assessed FOXO1 interaction with a known site within the PDK4 promoter region.^27,28^ High glucose exposure reduced the amount of FOXO1-bound to the PDK4 promoter compared to the normal glucose control (100±4 to 9±1 %, 91% decrease, *P*<0.0001, Fig. 9C). Hypoxia exposure significantly reduced FOXO1-bound PDK4 promoter compared to the normoxia control (100±4 to 39±5 %, 62% decrease, *P*<0.01). However, rHDL treatment under high glucose and hypoxic conditions enhanced the amount of FOXO1-bound PDK4 promoter sequence (34±3 to 61±13 %, 79% increase, *P*<0.05). Taken together, these findings suggest rHDL prevents the nuclear exclusion of FOXO1 (by reducing FOXO1 phosphorylation) and increases binding to the PDK4 promoter, presenting FOXO1 as a key transcription factor that regulates the increase in PDK4 by rHDL.

**Figure 9.**
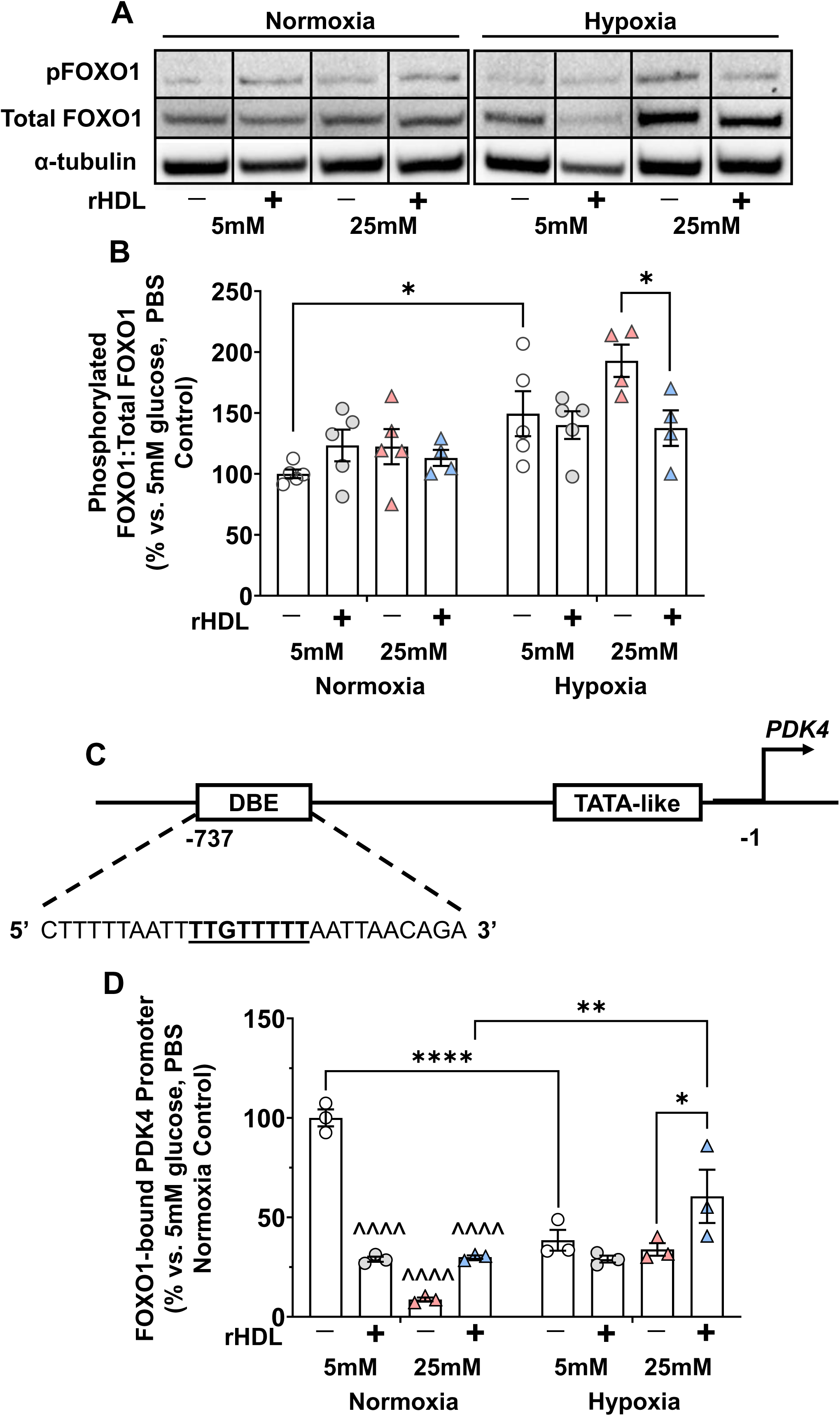
rHDL increases FOXO1 activity and binding to the PDK4 promoter in high glucose and hypoxia. HCAECs were incubated with rHDL (20 µM) or PBS (vehicle control) for 18 h, then exposed to normal (5 mmol/L) or high (25 mmol/L) glucose conditions for 72 h. Following this, cells were exposed to normoxia or hypoxia (1.2% O_2_) for 6 h. (A) Representative Western blot images of phosphorylated FOXO1 (pFOXO1) and total FOXO1 levels. (B) Graphed densitometry analyses of pFOXO1:Total FOXO1, with data normalized to α-tubulin (n=5). \**P*<0.05 by two-way ANOVA. (C) A schematic showing the FOXO1 binding site sequence (underlined) in the PDK4 gene promoter region that was targeted by the chromatin immunoprecipitation assay (ChIP). (D) FOXO1 binding to this known site in the PDK4 promoter was measured using ChIP (n=3). \**P*<0.05, \*\**P*<0.01, \*\*\*\**P*<0.0001 vs. controls by two-way ANOVA. ^^^^*P*<0.0001 vs 5mM glucose, PBS control by two-way ANOVA. 5 mM glucose, PBS (white circles), 5 mM glucose, rHDL (grey circles), 25 mM glucose, PBS (red triangles), 25 mM glucose, rHDL (blue triangles). Data are expressed as mean±SEM.

## DISCUSSION

Diabetes impairs ischemia-driven angiogenesis, a significant contributor to the development of diabetic vascular complications.^14^ EC metabolic reprogramming in response to hypoxia is an important part of preserving EC function. The PDK4/PDC axis is central to the regulation of this oxygen conserving mechanism,^13^ yet has never been implicated in the impairment of angiogenesis in diabetes. We have previously demonstrated that rHDL promotes physiological angiogenesis in response to ischemia in diabetic murine models.^14^ We now report the following important findings: (1) Diabetes impairs the PDK4/PDC axis and angiogenic responses to wound ischemia *in vivo* (2) rHDL rescues this, in parallel with a restoration of impaired wound neovascularization and healing. These findings were reiterated *in vitro*, where (3) high glucose impairs EC metabolic reprogramming responses to hypoxia, with rHDL incubation returning the regulation of the PDK4/PDC axis and mitochondrial respiration to that seen in normal glucose with; (4) restoration of high glucose-impaired *in vitro* angiogenic functions. Lentiviral shSR-BI knockdown experiments *in vitro* revealed the induction of PDK4 by rHDL is mediated by SR-BI. (5) Furthermore, *in vitro* and *in vivo* studies showed that inhibition of PDK4 ameliorates the proangiogenic effects of rHDL. Finally, (6) the mechanism underlying the metabolic effects of rHDL may be elicited via the activation of PDK4 by FOXO1 (Fig. 10). Taken together, we show diabetes impairs metabolic reprogramming responses to hypoxia and demonstrate a new mechanism by which rHDL rescues diabetes-impaired angiogenesis.

**Figure 10.**
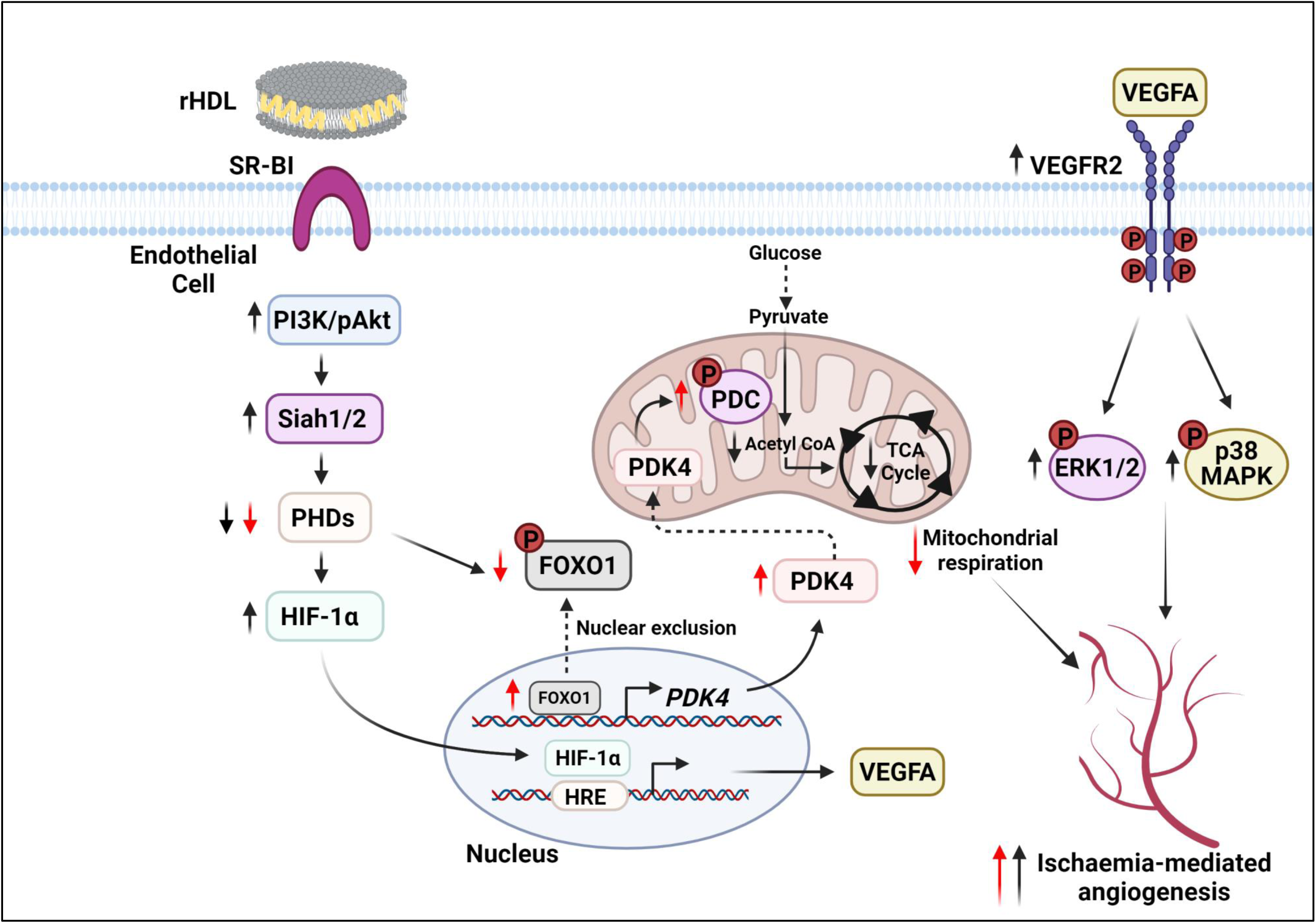
Proposed mechanism for the impairment of the PDK4/PDC axis response to ischemia by diabetes and its rescue by rHDL. We have previously shown that rHDL rescues diabetes-impaired angiogenesis by targeting multiple angiogenic signalling mediators.^14^ This includes the activation of the PI3K/Akt pathway, which induces the expression of the E3 ubiquitin ligases, Siah1 and Siah2. Increases in Siahs result in the inhibition of the prolyl hydroxylase domain (PHD) protein family. Suppression of PHD2/3 prevents HIF-1a from degradation, allowing it to translocate to the nucleus and bind to the hypoxia response element (HRE), activating transcription of proangiogenic mediators including VEGFA. VEGFA is released into the circulation, where it binds and phosphorylates VEGFR2, further augmenting angiogenesis by activating ERK1/2 and p38 MAPK signalling.^43^ We now show that rHDL rescue diabetes-impaired endothelial cell metabolic reprograming in hypoxia (denoted by red arrows). Diabetes impairs the induction of the PDK4/PDC axis in response to hypoxia, which is associated with an aberrant increase in mitochondrial respiration and an impairment to EC angiogenesis. rHDL decreases expression of the PHD proteins, which may affect phosphorylation of FOXO1. Decreased phosphorylation of FOXO1 leads to increased transcription factor activity and binding of FOXO1 to the PDK4 promoter region, which promotes the transcription of PDK4. Increased PDK4 is associated with an increase in inactivation (phosphorylation) of the PDC, which also occurs with rHDL treatment. Suppression of PDC activity decreases the amount of glucose-derived acetyl CoA available to fuel the TCA cycle and reduces mitochondrial respiration, which rHDL treatment returns to normal glucose levels. This improves cell survival in high glucose and hypoxia and provides an alternate pathway for the proangiogenic effects of rHDL in diabetes. These effects of rHDL are mediated via SR-BI.

We observed a striking impairment to the induction of PDK4 and pPDC in diabetic mice in the early stages post-induction of wound ischemia. While several studies have highlighted the importance of glycolysis in EC angiogenesis,^9,29^ few have examined suppression of mitochondrial respiration as an oxygen-conserving mechanism in hypoxia-driven angiogenesis. This result indicates that control of mitochondrial respiration is perturbed in diabetic wound tissue, and may contribute to impaired neovascularisation.

Previous studies have reported that diabetes modulates PDK4 expression, depending upon the tissue. For example, *PDK4* mRNA expression is elevated in muscle tissue from diabetic mice, indicating diabetes causes impaired glucose utilisation.^30^ Other studies have shown that general cellular metabolism is perturbed in different ways across diabetic tissues.^31^ This variation is expected, since different tissues display unique metabolic phenotypes that support their function. Our findings highlight the clear effect of diabetes on metabolic reprogramming in murine wound tissue, but also the importance of the PDK4/PDC axis in the early response to wound ischemia. A strength of this study is the measurement of the PDK4/PDC response *in vivo* over time post-wound ischemia in cohorts of mice. This provided a clear demonstration of the expression patterns in response to ischemia and revealed that the axis plays a lesser role in the later stages post-wound ischemia. The tissue-specific and temporal elements must both be considered in the translation of these findings.

rHDL enhanced PDK4 and pPDC levels in wound tissues of diabetic mice, and enhanced both neovascularization and healing, suggesting an association between the PDK4/PDC axis and neovascularization. rHDL also increased wound *Pdk4* and wound angiogenesis in diabetic female mice early post-wounding. The elevation in wound angiogenesis occurred at day1 in female versus day 3 in male mice, suggesting sexual dimorphism in the angiogenic responses to ischemia. This is a phenomenon that is increasingly being appreciated,^32^ with evidence for differences in endothelial cell functions and eNOS having been observed in hypoxia/ischemia between males and females.

Consistent with previous findings,^33–36^ our *in vitro* studies found that PLPC, a key component of rHDL, appeared to be the primary driver of *PDK4* induction. *PDK4* gene expression was also augmented in rHDL modified by glycation or oxidation. Studies have previously shown that PDK4 is elevated in response to oxidative stress and advanced glycation,^37–40^ therefore it is possible that this induction may be a survival response. The scavenger receptor SR-BI has been implicated in mediating several of the endothelial protective effects of HDL including migration, tubulogenesis, and re-endothelialization.^14,35,41,42^ Using siRNA deletion of SR-BI, we also found that SR-BI plays an important role in the ability of rHDL to induce *PDK4* mRNA levels. Furthermore, siRNA deletion of PDK4 *in vitro* inhibited tubulogenesis and *in vivo* PDK4 inhibition attenuated the ability of rHDL to promote wound closure in diabetic mice. These studies demonstrate the importance of PDK4 for the proangiogenic effects of rHDL, which are mediated via the receptor SR-BI.

We demonstrated that hypoxia exposure increased *PDK4* mRNA expression and pPDC levels, highlighting the importance of this mechanism in the EC hypoxia response. Importantly, exposure to high glucose in hypoxia did not elicit an additional stepwise induction in PDK4 or pPDC. This represents a high glucose-induced dysregulation in the PDK4/PDC response to hypoxia as there was a subsequent failure to adequately supress mitochondrial respiration. Furthermore, in parallel these conditions negatively impacted angiogenic functions in response to hypoxia, suggesting that aberrant metabolic responses to hypoxia contribute to the impairment of angiogenesis in high glucose. The ability to preserve oxygen consumption in ischemia appears essential to maintaining EC functions, including angiogenesis.

In the absence of an effect of rHDL on the hypoxia transcription factor HIF-1α, previously linked to PDK4,^10^ we investigated an alternate transcription factor, FOXO1, known to interact with the PDK4 promoter region.^25^ When FOXO1 is phosphorylated, it is excluded from the nucleus and its transcriptional activity is abrogated.^25,27,28^ A role for FOXO1 in angiogenesis and EC function has also previously been identified. Wilhelm *et al*. demonstrated that deletion of FOXO1 in mice caused an increase in vessel sprouting.^26^ Conversely, its overexpression severely restricted angiogenesis and led to vessel thinning. We observed that in hypoxia with high glucose exposure, FOXO1 phosphorylation levels were at their highest, this increases nuclear exclusion and reduces FOXO1 transcription factor activity. Consistent with this finding, we found in these same conditions an increase in the binding of FOXO1 to the promoter region of PDK4 using ChIP. Furthermore, PHD3 levels were reduced by rHDL in high glucose and hypoxia. Interestingly, PHD3 has been demonstrated to maintain a regulatory relationship with FOXO1 phosphorylation.^24^ Taken together, inhibition of FOXO1 phosphorylation by rHDL presents as a mechanism to explain our observed induction of PDK4 by rHDL (Fig. 10).

In conclusion, we have demonstrated that diabetes impairs metabolic reprogramming and angiogenic responses to hypoxia, which are rescued by rHDL *in vitro* and *in vivo*. These findings provide further mechanistic support for the effects of rHDL on diabetes-impaired angiogenesis and has significant implications for the translation of HDL-based therapies, with the novel application of improving wound healing in diabetes.

## Acknowledgements

Not applicable

